# Neural Network Informed Photon Filtering Reduces Artifacts in Fluorescence Correlation Spectroscopy Data

**DOI:** 10.1101/2023.08.24.554627

**Authors:** Alva Seltmann, Pablo Carravilla, Katharina Reglinski, Christian Eggeling, Dominic Waithe

**Affiliations:** Institute for Applied Optics and Biophysics, Friedrich Schiller University Jena, Jena, 07743, Germany; Leibniz Institute of Photonic Technology, Jena, 07745, Germany; Science for Life Laboratory, Department of Women’s and Children’s Health, Karolinska Institutet, Stockholm, 171 77, Sweden; Jena University Hospital, Friedrich Schiller University Jena, Jena, 07747, Germany; MRC Weatherall Institute of Molecular Medicine, Radcliffe Department of Medicine, University of Oxford, Oxford, OX1 2JD, United Kingdom; work carried out whilst at: MRC Centre for Computational Biology and Wolfson Imaging Centre, MRC Weatherall Institute of Molecular Medicine, Radcliffe Department of Medicine, University of Oxford, Oxford, OX1 2JD, United Kingdom

**Keywords:** FCS, peak detection, super-resolution, artifact correction, deep learning, u-net

## Abstract

Fluorescence Correlation Spectroscopy (FCS) techniques are well-established tools to investigate molecular dynamics in confocal and super-resolution microscopy. In practice, users often need to handle a variety of sample or hardware-related artifacts, an example being peak artifacts created by bright, slow-moving clusters. Approaches to address peak artifacts exist, but measurements suffering from severe artifacts are typically non-analyzable. Here, we trained a 1-dimensional U-Net to automatically identify peak artifacts in fluorescence time-series and then analyzed the purified, non-artifactual fluctuations by time-series editing. We show that in samples with peak artifacts, the transit time and particle number distributions can be restored in simulations and validated the approach in two independent biological experiments. We propose that it is adaptable for other FCS artifacts, such as detector dropout, membrane movement, or photobleaching. In conclusion, this simulation-based, automated, open-source pipeline makes measurements analyzable which previously had to be discarded and extends every FCS user’s experimental toolbox.

## Introduction and Background

Fluorescence Correlation Spectroscopy (FCS) and its related techniques are powerful tools to unravel nanoscopic processes with applications in physics, chemistry, biology, and medicine.^1–3^ In their fifty-year-long history, these techniques have been used to examine photophysical and biomolecular dynamics on the scale of picoseconds to seconds and picomolar to micromolar molecular concentrations.^4–8^ The wide range of applications goes from analysing molecular interactions to gathering structural information on membranes, gaining insight into cellular processes and even whole organisms and tissues. ^9–15^ With increasing stability and streamlining of FCS methods, they are being deployed in high-throughput, quantitative settings. ^16–20^

The success of FCS is grounded in its core idea and its continuing technical improvement over the years. The core idea is to measure fluorescence fluctuations in a system in equilibrium over time. The experimental setups vary depending on the technique but usually include laser illumination, small probe volumes, fast photon detectors, and fast data processors. These fluctuations stem from many different processes, such as translational diffusion due to Brownian motion or photophysical properties such as triplet dynamics. By autocorrelating the fluorescence time-series and fitting the autocorrelation curves with theoretical models, researchers extract the desired information of transit times, triplet times, concentrations, or brightness.^4,21,22^ Importantly, if the system is not in equilibrium and these fluctuations are not ‘stationary’ around a mean value, FCS analyses may yield artifactual values. We refer to the existing literature for a general breakdown of FCS artifacts.^10,23–29^

Peak artifacts are common in biological FCS measurements and endanger the ‘condition of stationarity’. In challenging environments such as cells, researchers might find bright, slow-moving clusters stemming from internalized vesicles, cell debris, or unspecific binding of fluorophores.^13,28,30–34^ Other problematic settings are very long measurement times which make passing aggregates more likely^13,18,34–36^, movements of (sub) cellular components such as membranes^37–40^, or fastly aggregating probes such as up-conversion nanoparticles.^41^ Even single peak artifacts in the time-series, so-called rare events, bias the autocorrelation curve because each particle is weighted by the square of the particle brightness.^21,29^

In a typical FCS analysis workflow, the user visually inspects the autocorrelation function and underlying fluorescence time-series to ensure no artifacts are present. If peak artifacts arise, most common guides advise users to first adjust the biological sample and microscope parameters to avoid intensity shifts. Users might employ biophysical techniques such as prebleaching to bleach slow or immobile large clusters^30,42^ or biochemical techniques such as centrifugation to filter out aggregates.^26,43^ If peak artifacts persist, the most common advice is to hand-select time-series for further analysis and discard or manually crop time-series with peak artifacts.^10,13,26,29,30,34,44^ Additionally, broad peak artifacts might be correctable by methods intended to handle slower time-series deviations, such as photobleaching. These include dividing experiments into short intervals and averaging the resulting autocorrelation curves ^35,45^, purposefully avoiding the correlation of long lag times^17,46^, or adding a 1st order bias factor to the correlation function. ^47^ Lastly, users might improve fit outcomes by including an additional mobile species in the FCS fit equation^30^ and by weighting the autocorrelation function by its standard deviation when evaluating the goodness of fit.^48^

Typical biases arise because hand-selection is frequently ambiguous, fitting an additional mobile species often fails to capture the distorted parts of the correlation curve, and shortening the time-series can introduce a bias in the correlation function. ^47,49,50^ To circumvent these biases, Persson et al. introduced modulation filtering to remove peak artifacts with a rectangular intensity modulation of the fluorescence signal.^49^ A necessary premise for modulation filtering is a reliable segmentation of the time-series to find these artifacts. Furthermore, Baum et al. proposed Fourier filtering to filter out slow components of the fluorescence signal^39^, although an independent study found only incomplete artifact removal.^40^

Lastly, multiple automation approaches aim to remove the subjective bias and ambiguity of hand-selection and enable FCS for high-throughput settings. ^50–52^ As exemplified by Ries et al., such an algorithm splits one time-series into fixed parts and correlates them, removes correlations deviating too far from the average, and averages the rest. Limitations are that a user still has to choose algorithmic parameters manually, such as the time window for splitting, or the threshold for removal. Also, the correlation bias for short time-series can still be an issue.

New deep learning-based techniques promise to further reduce the number of subjective decisions of the user during analysis. Neural networks learn to identify patterns in data beyond the capabilities of conventional signal processing strategies. In microscopy, they excel at computer vision tasks such as classification, object tracking, and segmentation. ^53^ In FCS techniques specifically, Ren used a neural network to directly predict diffusion coefficients from imaging FCS autocorrelation curves. ^54,55^ Even though the potential of neural networks is eminent, a crucial point for most neural network projects is gathering training data.

In this paper, we first describe simulation experiments on how to deal with peak artifacts in FCS time-series. Second, we report a neural network for peak artifact segmentation. Third, we validate the neural network and an automated, open-source correction pipeline in two applied experiments. This approach enables photon filtering based on time-series segmentation of artifacts visible in the fluorescence signal.

## Results and discussion

As highlighted, we in a first step discuss and compare correction methods, meaning automated approaches for refining FCS time-series to ensure more dependable FCS analyses. Further, we discuss and compare segmentation methods, which offer FCS time-series segmentations as a foundation for the correction methods. Finally, we assess an automated pipeline that combines segmentation and correction techniques on separate datasets.

### Correction methods and simulations

In this study, the term ‘correction methods’ describes approaches to improve FCS analyses suffering from visible artifacts in the fluorescence timeseries. These optimization approaches had to integrate into a fully automated pipeline without manual parameter tuning. They also had to work on FCS time-series and time-correlated single photon counting (TCSPC) data. As illustrated in Figure 1, we chose three methods: *cut and stitch* ^32,49^, *set to zero* (inspired by statistical filtering in FCS^56,57^), and *averaging* .^35,45^ The input for each method was a binned intensity time-series (for simulation experiments) or TCSPC data (for applied experiments) and a segmentation vector marking parts with and without peak artifacts. *Cut and stitch* meant cutting out bins with peak artifacts, stitching the remaining ones together, and correlating the resulting time-series. *Set to zero* meant setting the intensity value of each bin with peak artifacts to zero before correlating the resulting time-series. *Averaging* meant correlating each instance of an intact segment without peak artifacts and averaging the autocorrelation curves. In summary, the correction methods only changed the intensity time-series (*cut and stitch, set to zero*) or the correlation (*averaging*), but not the FCS fit.

**Figure 1.**
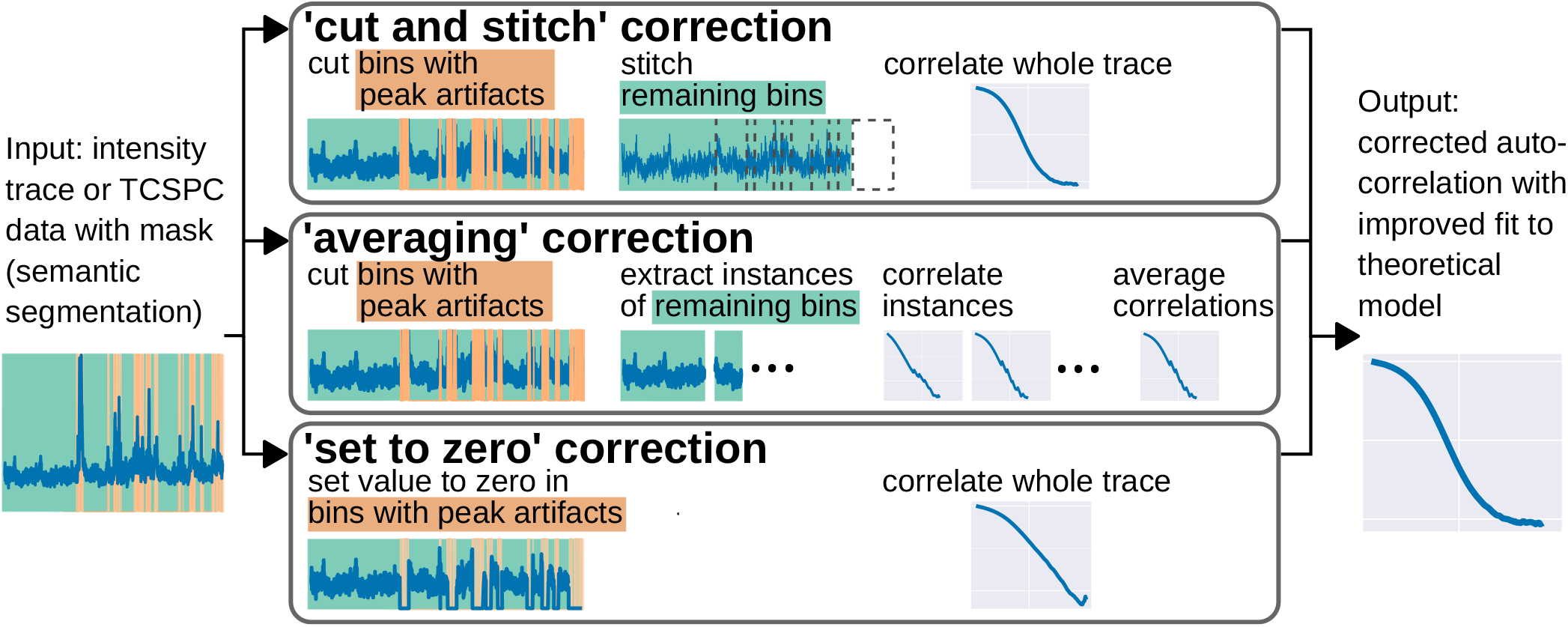
Principles of correction methods. *Set to zero* preserves time step order but introduces values far from the average, which can dominate the correlation. *Averaging* correlates instances without artifacts and preserves time step order but suffers from FCS bias and noise due to the shortened time-series. *Cut and stitch* (also called *gap contraction* ^49^) does not suffer from FCS bias and noise and does not introduce values far from the average but introduces ‘stitching artifacts’ where the time-series parts are stitched together. For a further analysis of ‘stitching artifacts’ see Figure S1.

To test these correction methods, and later train artifact prediction models, we produced various FCS time-series using 2D Monte Carlo simulations of particles. These simulations yielded three corresponding vectors per time-series: intensity over time without artifacts, intensity over time with peak artifacts, and a segmentation vector showing the location of the artifacts. We achieved diverse time-series shapes by simulating molecules (low brightness) and clusters (high brightness) with different transit times through the 2-dimensional observation spot. The molecule transit time *τ*_mol_ determined time-series segments without artifacts, and the cluster transit time *τ*_clust_ determined time-series segments with artifacts, as well as the segmentation vector. Further details on the simulations are described in Supplementary Note.

Figure 2A shows a subset of 9 subgroups of simulations that we used to test the correction methods. Our metric for a successful correction was a close agreement of the FCS fit outcomes after peak artifact correction compared to FCS fit outcomes of time-series without artifacts. ‘Fit outcomes’ refers to either the transit time through the observation spot *τ*_*D*_ or the particle number *N*. When analysing a large number of measurements without artifacts, *N* shows a normal distribution and *τ*_*D*_ shows a log-normal distribution.^18^ The simulations illustrated a strong shift in the *τ*_*D*_ distribution between time-series with and without artifacts for three different simulated *τ*_mol_ (Figure 2B). The three correction methods were applied to time-series with peak artifacts and artifacts were identified using the simulated segmentation vectors. *Cut and stitch* restored transit time distributions independently of *τ*_mol_, while *averaging* and *set to zero* only showed slight improvements for *τ*_mol_ = 163.35 ms. When comparing illustrative correlation and fit curves (see Figure S2), *cut and stitch* restored autocorrelations curves to be similar to curves from data without peak artifacts, while *averaging* and *set to zero* showed strong deviations.

**Figure 2.**
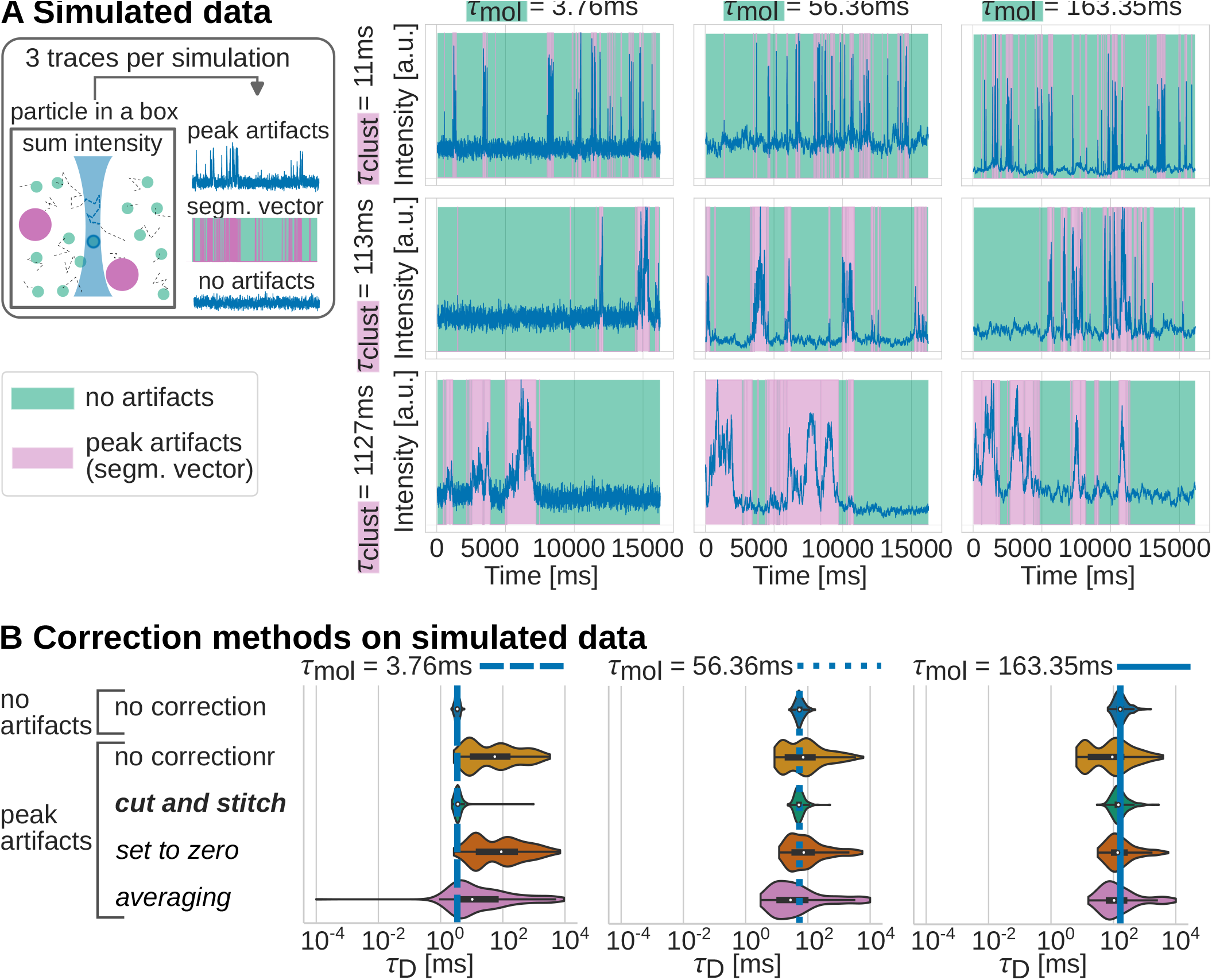
Simulated data for evaluation of correction methods and model training. **A** Sketch of simulation process and illustrative plots of 9 subgroups of simulated data used for testing the correction methods. In total, we used 30 groups for U-Net training and evaluation. The molecule transit time *τ*_mol_ determined segments of the time-series without artifacts (green), and the cluster transit time *τ*_clust_ determined segments of the time-series with artifacts and the segmentation vector (pink). **B** Comparing correction methods on batches of simulated data with peak artifacts by evaluating transit time *τ*_*D*_ distributions from FCS correlation and fit. Expected transit times *τ*_mol_ and segmentation vectors for the correction were extracted from simulations. All groups (left labels) with peak artifafcts yielded three distributions (1 species fit, fast sp. of 2 sp. fit, slow sp. of 2 sp. fit). Here, we only displayed the distribution closest to the expected value. For illustrative correlation and fit curves, extended fit outcomes, and simulation details see Figures S2 and S3A and Supplementary Note.

The overall good performance of *cut and stitch* confirms prior results by Persson et al.. This result might be counter-intuitive because this method does not keep the time axis intact. To test the limits of *cut and stitch*, we investigated the effect of randomly placed cuts on time-series without artifacts (see Figure S1). The plot shows that introducing a certain number of cuts is possible without shifting the fit outcome distributions, depending on transit time, time-series length, and time resolution. Our simulation methods enable users interested in the *cut and stitch* approach to employ similar control plots for their time-series parameters of interest. Furthermore, we attribute the low performance of *averaging* to lower signal-to-bias and signal-to-noise ratios stemming from shortened time-series. The correlation curve bias refers to a systematic deviation for short time-series compared to a theoretical curve calculated from infinite experiment time. ^47^ Correlation curve averaging can only reduce the noise but not the bias. This effect of shortened time-series also affects *cut and stitch*, although we only observed a slight broadening of the distributions and no apparent bias. Lastly, the low performance of *set to zero* stems from introducing time-series values, which are very far from the average and dominate the correlation function. These unwanted correlations are stronger the higher the average number of photon counts is.

### Peak artifact segmentation

To automatically predict peak artifacts in FCS time-series, we employed the U-Net deep learning network.^58,59^ U-Net architectures are widely used when tackling biomedical or cell segmentation tasks and are one of the state-of-the-art methods for this task in supervised machine learning.^53,60–62^ As shown in Figure 3A, our implementation was a standard 1-dimensional U-Net with an added padding step, which allowed segmenting time-series of arbitrary length. U-Nets are suitable for situations requiring an output of similar dimension and resolution to our input. In this case, we classified the presence or absence of artifacts across our signal at a fine-grain resolution, making a U-Net a highly suitable choice. We trained the model on 30 subgroups of simulated data (Figure 2 displays 9 of these subgroups), thus enabling the network to generalize well to slower and faster dynamics in the time-series and to identify broader and more narrow peak artifacts. We chose the best model hyperparameters on a simulated validation dataset and evaluated the prediction performance on a simulated, held-out test dataset. By applying well-performing models in the automated correction pipeline described later, we validated their usefulness on an independent dataset while using fit outcomes as an independent metric. In these applied experiments, we also compared the U-Net predictor with a manually chosen threshold after robust scaling (see Figure 3B). For details about model training and evaluation, see the Supplementary Note and Tables S1 and S2.

**Figure 3.**
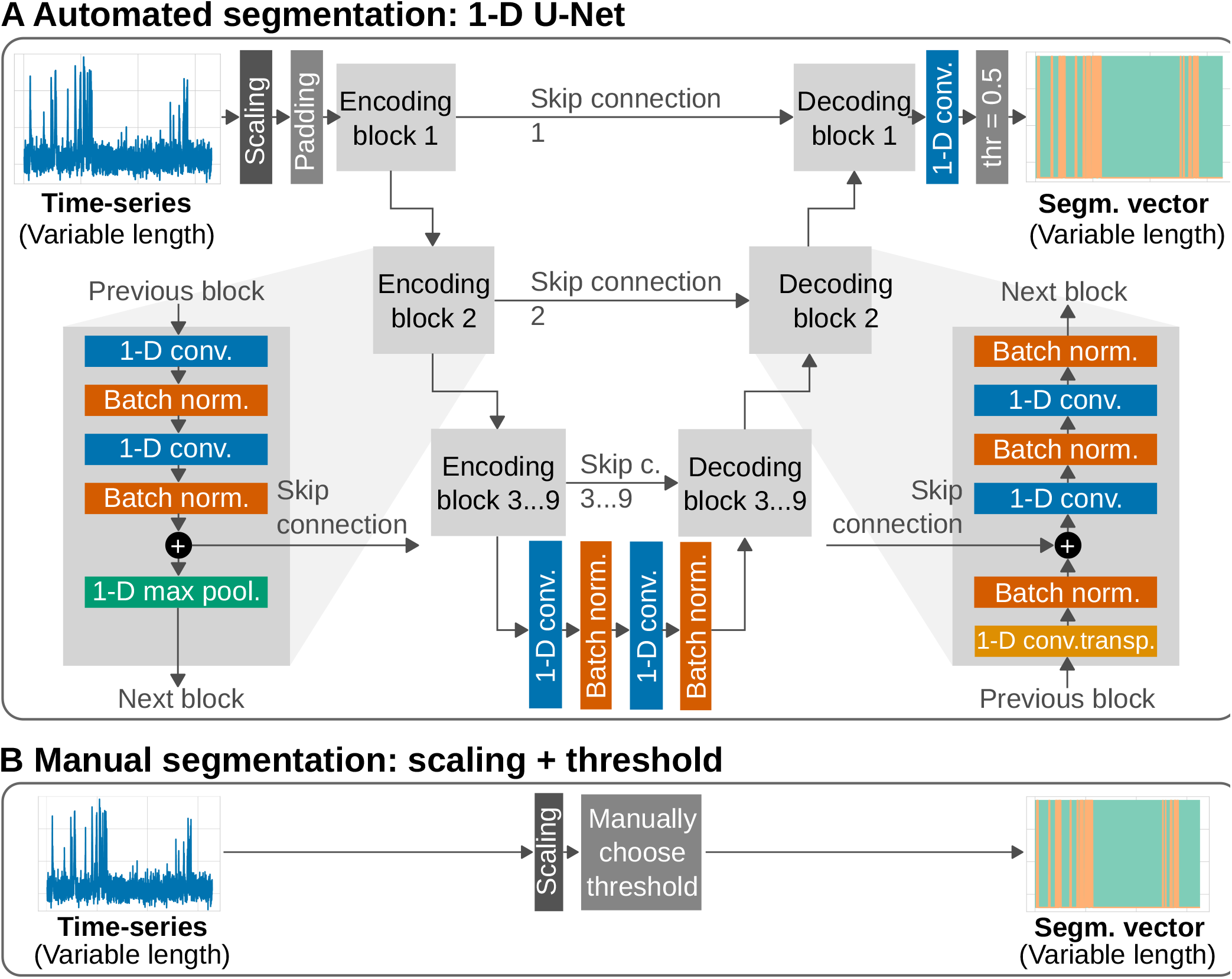
Principles of segmentation methods. **A** Architecture of the fully convolutional 1-dimensional U-Net used in the automated pipeline. The padding step allowed a variable input length. **B** Manual segmentation by robust scaling and a threshold chosen by trial and error. For further details see the Supplementary Note and Tables S1 and S2.

These results add to existing research on using U-Nets in 1-dimensional signal processing. Examples are the segmentation of electrocardiograms^63–65^, segmentation of vesicle trajectories^66^, and Raman and infrared spectra refinement.^67,68^ To our knowledge, this is the first application of U-Nets on FCS intensity time-series.

### Applied experiments and automated pipeline

The following analysis validated the U-Net segmentation method and established a usable, automated pipeline to reduce subjective decisions in FCS analyses. First, we describe two applied experiments yielding FCS measurements with peak artifacts and a biologically similar control without peak artifacts. In the first applied experiment (Figure 4A), we compared the small dye Alexa Fluor 488 (AF488, 643.4 Da^69^) in solution and a mix of AF488 and large unilamellar vesicles labelled with the dye DiO (DiO-LUVs). These artificial lipid vesicles had a diameter of around 100 nm and contained multiple DiO molecules. They were several orders of magnitude slower and brighter than the diffusing fluorophore AF488 and produced peak artifacts with the steady fluctuation of AF488 in the background. The second applied experiment (Figure 5A) examined two homologues of the peroxisomal import receptor protein PEX5 fused with eGFP, Trypanosoma brucei-PEX5-eGFP (Tb-PEX5-eGFP) and Homo sapiens-PEX5-eGFP (Hs-PEX5-eGFP). For unknown reasons, Tb-PEX5-eGFP partially formed aggregates in solution, leading to a steadily fluctuating signal with intermittent peak artifacts. The homologue from another species, Hs-PEX5-eGFP, only showed a steadily fluctuating signal. Both molecules had a similar structure and molecular weight, making their transit times from FCS fitting comparable. For further details on the applied experiments see the Supplementary Note.

We compared the correction methods while keeping the U-Net segmentation method fixed (Figures 4B and 5B) and the segmentation methods while keeping the *cut and stitch* correction method fixed (Figures 4C and 5C). These plots show the distributions of the main fit outcome parameters *τ*_*D*_ and *N* for large batches of timeseries, as detailed in the Supplementary Note.

For AF488 experiments, the median *τ*_*D*_ (IQR) and *N* (IQR) for time-series without artifacts was 39.7 µs (39.0–40.4) and 14.3 (14.2–14.5), and with artifacts was 59.7 µs (42.5–117.8) and 0.9 (0.5–1.5). Figure 4B shows that the automated pipeline with *cut and stitch* best restored the fit outcomes, with a median *τ*_*D*_ (IQR) of 38.8 µs (35.2–42.2) and *N* (IQR) of 15.2 (14.3–16.0). The other correction methods were not sufficient, with *set to zero* yielding 102.0 µs (73.2–151.8) and 0.5 (0.4–0.6) and *averaging* yielding 82.4 µs (69.4–96.0) and 18.1 (17.0–19.1). Figure 4C shows that the automated pipeline was non-inferior to the manual pipeline, which yielded 43.3 µs (40.6–46.8) and 12.0 (10.3–13.5). Also, neither method introduced severe shifts in the distribution when applied to time-series without artifacts, with the automated pipeline yielding 39.6 µs (38.9–40.3) and 14.3 (14.2–14.5) and the manual pipeline yielding 38.1 µs (37.4–38.8) and 14.5 (14.3–14.6). The slight *τ*_*D*_ shift in the manual pipeline shows that the simple decision boundary in thresholding inevitably will segment some normal fluctuation peaks as peak artifacts. In the best case, this effect is not present in the learned decision boundary of the automated pipeline.

**Figure 4.**
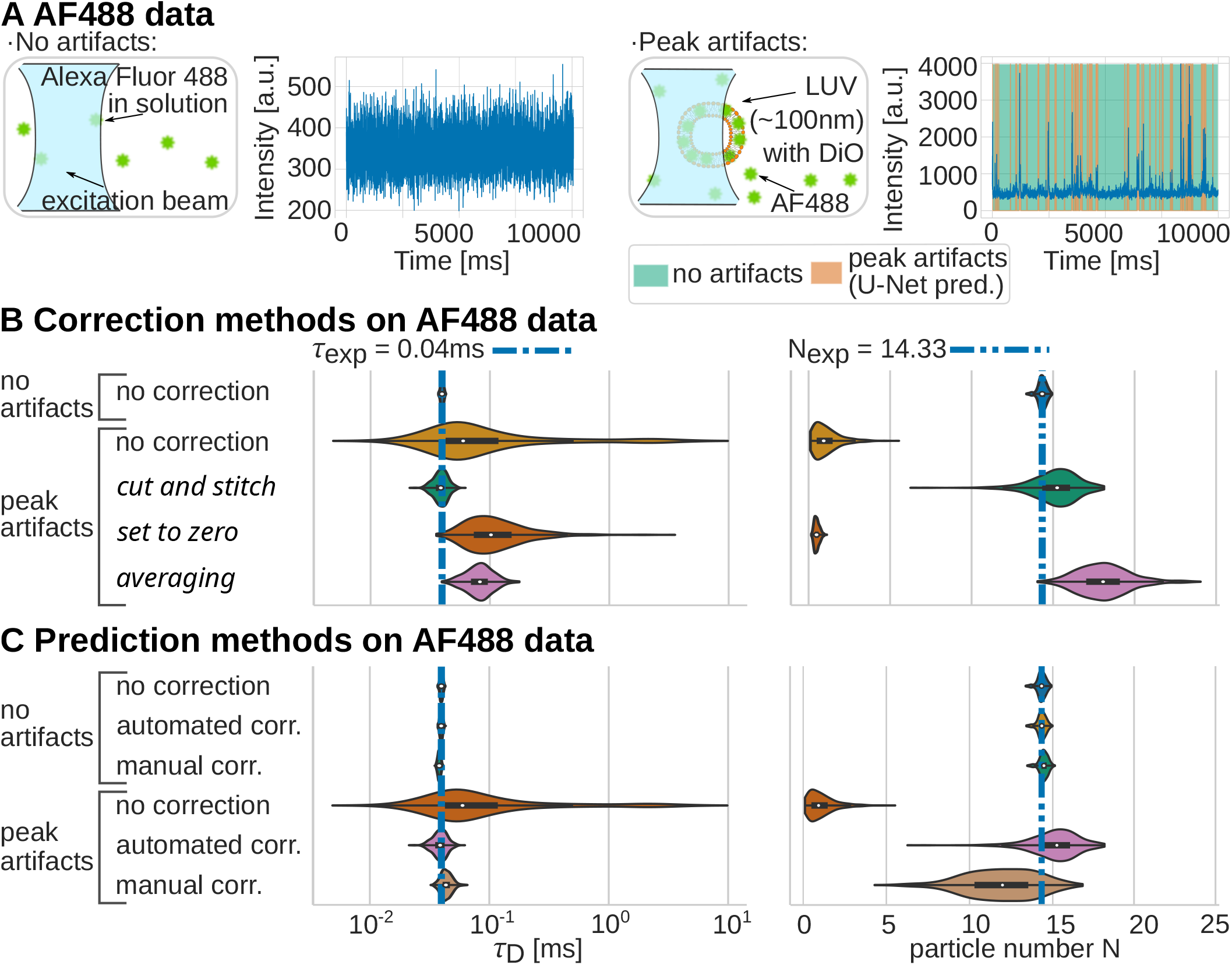
Applied AF488 experiments. **A** Sketches (not to scale) and illustrative plots of AF488 data with and without peak artifacts. Segmentation by U-Net model (orange). **B-C** Evaluation of transit time *τ*_*D*_ and *N* distributions from FCS correlation and fit for different correction and segmentation methods. The automated pipeline of U-Net predictor and *cut and stitch* successfully restored both distributions. Expected transit times *τ*_exp_ and particle numbers *N*_exp_ from the group ‘No artifacts, no correction’ (median of 1 component fits). For further details see the Supplementary Note and Figures S3B and S4A.

For PEX5 experiments, the median *τ*_*D*_ (IQR) for time-series without artifacts was 356 µs (346– 368) and with artifacts was 437 µs (404–498). Figure 5B shows that all correction methods improved the fit outcome distribution, but *cut and stitch* slightly over-corrected the results. In detail, the automated pipeline with *cut and stitch* yielded 325 µs (309–343), *set to zero* yielded 366 µs (348–383), and *averaging* yielded 349 µs (349–349). Figure 5C shows that the manual pipeline correctly restored the *τ*_*D*_ distributions to 362 µs (344–383) and did not severely shift the distribution when applied to time-series without artifacts, yielding 348 µs (327–360). On the contrary, the automated pipeline introduced a shift when applied to time-series without artifacts, yielding 324 µs (308–336). The worse performance of the automated pipeline in PEX5 data in both time-series with and without artifacts points to a phenomenon known in the machine learning field as ‘covariate shift’.^70^ It describes diminished model performance if the training data did not capture the features of the application data. One such example was that the mean intensity between simulated training data (500– 3500 a.u.) did not include the mean intensity of PEX5 data (about 50 a.u.). This lower average intensity also explains the better performance of the *set to zero* correction method, which for PEX5 data did not introduce time-series values far from the average.

**Figure 5.**
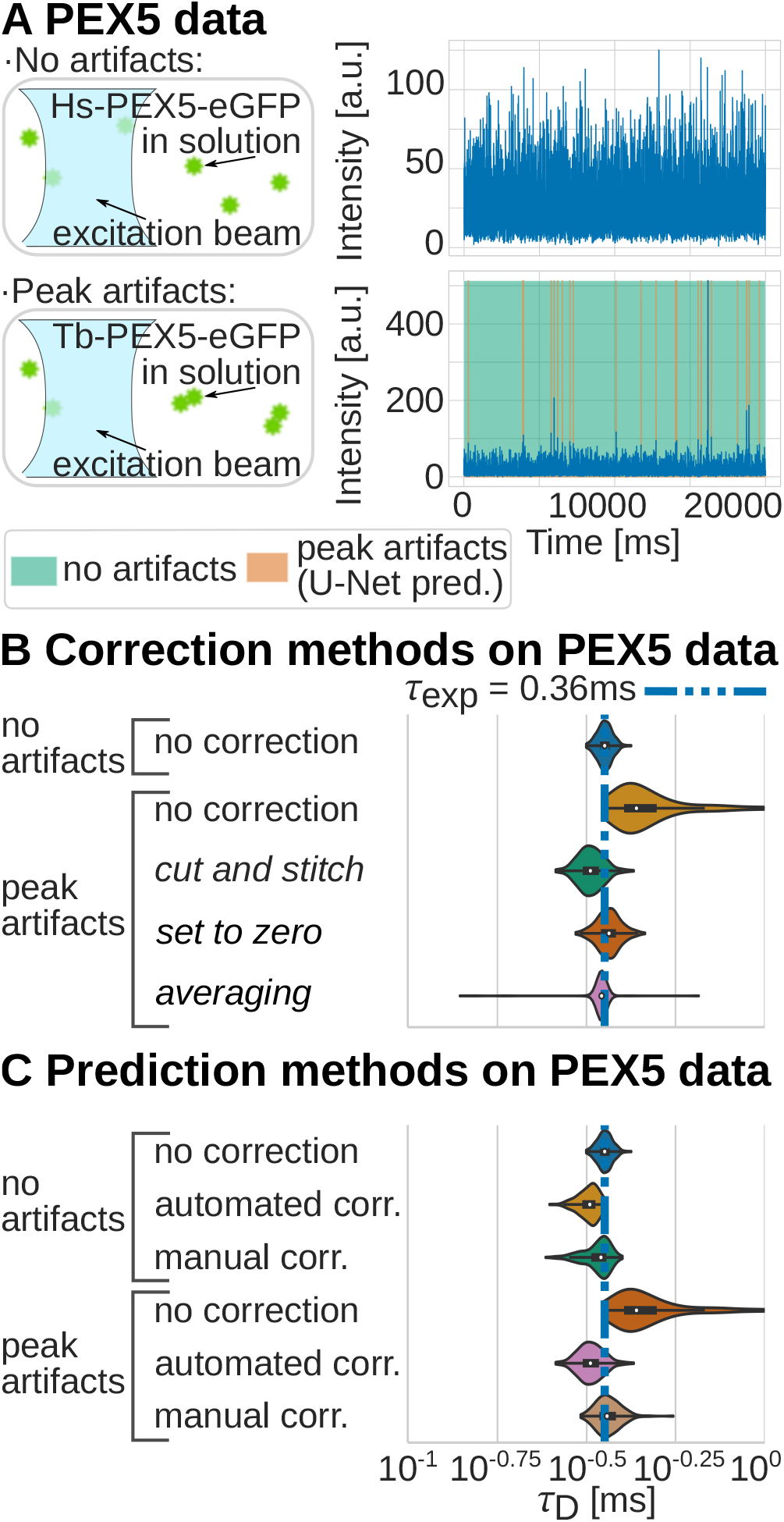
Applied PEX5 experiments. **A** Sketches (not to scale) and illustrative plots of PEX5 data. Segmentation by U-Net (orange). **B-C** Evaluation of transit time *τ*_*D*_ distributions from FCS correlation and fit analogous to Figure4. Expected transit time *τ*_exp_ from the group ‘No artifacts, no correction’ (median of 1 component fits). Experiments and FCS fit as described in the Supplementary Note and Figures S3C and S4B.

Users interested in the automated pipeline should be aware of the ‘covariate shift’ effect, which can usually be mitigated by simulating data resembling the intensity profiles of PEX5 data and re-training the network. Machine learning-specific quality controls are necessary if users re-train a model, as described elsewhere.^53,60^ Lastly, users should have a good understanding of their biological sample. Peak artifacts stemming from stained vesicles, dye aggregations, or membrane movement occur regularly and interfere with the desired signal. But peak artifacts might be biologically relevant, as in FCS studies on protein aggregation.^71,72^

## Conclusions

We presented an automated, open-source pipeline to optimize FCS data with unwanted peak artifacts. The primer for our corrective approach was a simulation method which allowed training our U-Net neural network without manually annotating training data. As a key piece, this network successfully predicted peak artifacts in fluorescence time-series and was independently evaluated in two applied experiments. Furthermore, we showed that the *cut and stitch* approach is an excellent method for reconstructing the time-series, improving the outcome of the subsequent correlation and fit. Although this study used point FCS data, these approaches apply to scanning FCS, imaging FCS, and other FCS varieties, if the researcher has access to photon time-series. Researchers interested in high-throughput applications will find the automated pipeline valuable as well. All in all, this toolbox enables FCS measurements on data which users previously would have discarded.

To make applying the presented methods as accessible as possible, we provide the low-code Google Colab notebook ‘U-Net for FCS’ at https://github.com/HenriquesLab/ZeroCostDL4Mic. As part of the Zero-CostDL4Mic toolbox^60^, it enables FCS time-series simulation, model (re-)training and evaluation, and application of the automated or manual pipeline on TCSPC data. Furthermore, we provide a version-controlled record of all data, code, and analysis decisions at https://github.com/aseltmann/fluotracify. Future work can extend this toolbox in multiple ways. The U-Net predictor can be adapted for multi-class semantic segmentation to discriminate between various time-series artifacts, such as peak artifacts, photobleaching artifacts, membrane movements, and detector dropout artifacts. Instead of optimizing FCS fit outcomes, users can employ the segmentation vector to gain insight of time-series segments showing FCS artifacts, which might be desirable in protein aggregation analysis. Also, alternative correction regimes such as modulation filtering can substitute the pragmatic *cut and stitch* approach and make the pipeline more robust. We believe that extending open-source, easy-to-use, automated tools will facilitate new and improved FCS measurements in challenging or high-throughput experimental settings and provide reliable results to researchers investigating the nanoscale.

## Acknowledgement

The authors thank Falk Schneider, Stefan Hoffmann, and Thilo Figge for discussions and feedback during early stages of the project. This research was supported by the UK Research and Innovation Biotechnology and Biological Sciences Research Council (BB/P026354/1) and the UK Research and Innovation Medical Research Council (MR/S005382/1a, MC UU 12009, MC UU 12010 / unit programs G0902418 and MC UU 12025, MR/K01577X/1). The computational experiments were performed on resources of FSU Jena supported in part by German Research Foundation (DFG) grants INST 275/3341 FUGG and INST 275/363-1 FUGG. P.C. received funding from the European Commission Horizon 2020 Marie Sk1odowska Curie Programme (H2020-MSCA-IF-2019-ST project 892232 FILM-HIV) and the Basque Government (POS 2018 1 0066 and POS 2019 2 0022). K.R. received funding from the DFG (research unit 1905, grant no. 322325142). C.E. further acknowledges financial support by the DFG (Germany’s Excellence Strategy – EXC 2051 – Project-ID 390713860, and project number 316213987 – SFB 1278), the Free State of Thuringia (TMWWDG; TAB; AdvancedSTED / FGZ: 2018 FGI 0022; Advanced Flu-Spec / FGZ: 2020 FGI 0031), the innovation program by the German Federal Ministry for Economic Affairs and Climate Action (BMWK; ZIM -project 16KN070934 / Lab-on-a-chip FCS-Easy), the German Federal Ministry for Education and Research (BMBF; funding program LIVE2QMIC -FGZ: 13N15956, as well as Photonics Research Germany -FKZ: 13N15713 / 13N15717). This work is integrated into the Leibniz Center for Photonics in Infection Research (LPI). The LPI initiated by Leibniz-IPHT, Leibniz-HKI, UKJ and FSU Jena is part of the BMBF national roadmap for research infrastructures.

## Supporting Information Available

Supplementary Note detailing experimental methods and materials, Supplementary Tables on supervised machine learning and FCS fit equations, Supplementary Figures detailing extended FCS fit outcomes, stitching artifacts, and illustrative correlations and fit curves.

## TOC Graphic

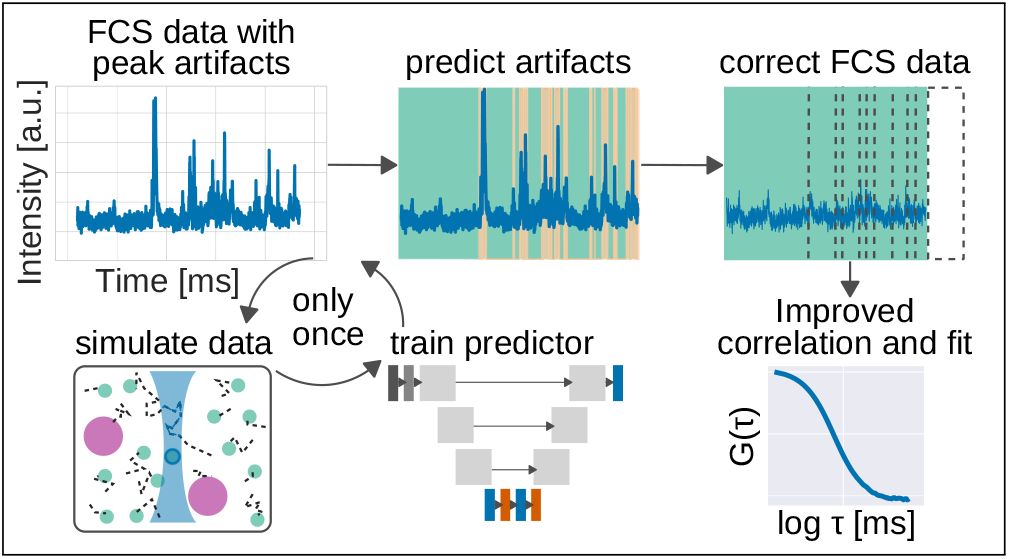

## Notes

https://github.com/aseltmann/fluotracify

https://doi.org/10.5281/zenodo.8137129

https://doi.org/10.5281/zenodo.8074408

https://doi.org/10.5281/zenodo.8082558

https://doi.org/10.5281/zenodo.8109282

### Competing Interest Statement

The authors have declared no competing interest.

## References

(1) Magde, D.; Elson, E.; Webb, W. W. Thermodynamic Fluctuations in a Reacting System—Measurement by Fluorescence Correlation Spectroscopy. Physical Review Letters 1972, 29, 705–708, DOI: 10.1103/PhysRevLett.29.705.

(2) Ehrenberg, M.; Rigler, R. Rotational brownian motion and fluorescence intensify fluctuations. Chemical Physics 1974, 4, 390–401, DOI: 10.1016/0301-0104(74)85005-6.

(3) Eggeling, C.; Hellriegel, C. Editorial. Methods 2018, 140-141, 1–2, DOI: 10.1016/j.ymeth.2018.04.034.

(4) Wohland, T.; Maiti, S.; Macháň, R. An Introduction to Fluorescence Correlation Spectroscopy ; IOP Publishing, 2020.

(5) Sasaki, A.; Halter, M.; Elliott, J. T. Fluorescence Correlation Methods for Determining Absolute Numbers of Molecules from Microscopy Images. bioimages 2019, 27, 13–22, DOI: 10.11169/bioimages.27.13.

(6) Fitzpatrick, J. A.; Lillemeier, B. F. Fluorescence correlation spectroscopy: linking molecular dynamics to biological function in vitro and in situ. Current Opinion in Structural Biology 2011, 21, 650–660, DOI: 10.1016/j.sbi.2011.06.006.

(7) Widengren, J.; Mets, U.; Rigler, R. Fluorescence correlation spectroscopy of triplet states in solution: a theoretical and experimental study. The Journal of Physical Chemistry 1995, 99, 13368–13379, DOI: 10.1021/j100036a009.

(8) Eggeling, C.; Widengren, J.; Rigler, R.; Seidel, C. A. M. Photobleaching of Fluorescent Dyes under Conditions Used for Single-Molecule Detection: Evidence of Two-Step Photolysis. Analytical Chemistry 1998, 70, 2651–2659, DOI: 10.1021/ac980027p.

(9) Rigler, R.; Elson, E. S. In Fluorescence Correlation Spectroscopy: Theory and Applications; Schäfer, F. P., Toennies, J. P., Zinth, W., Eds.; Springer Series in Chemical Physics; Springer Berlin Heidelberg: Berlin, Heidelberg, 2001; Vol. 65; DOI: 10.1007/978-3-642-59542-4.

(10) Sezgin, E.; Schwille, P. Fluorescence Techniques to Study Lipid Dynamics. Cold Spring Harbor Perspectives in Biology 2011, 3, 32, DOI: 10.1101/cshperspect.a009803.

(11) Elson, E. Fluorescence Correlation Spectroscopy: Past, Present, Future. Biophysical Journal 2011, 101, 2855–2870, DOI: 10.1016/j.bpj.2011.11.012.

(12) Macháň, R.; Wohland, T. Recent applications of fluorescence correlation spectroscopy in live systems. FEBS Letters 2014, 588, 3571–3584, DOI: 10.1016/j.febslet.2014.03.056.

(13) Sezgin, E.; Schneider, F.; Galiani, S.; Urbaňcič, I.; Waithe, D.; Lagerholm, B. C.; Eggeling, C. Measuring nanoscale diffusion dynamics in cellular membranes with super-resolution STED–FCS. Nature Protocols 2019, DOI: 10.1038/s41596-019-0127-9.

(14) Gupta, A.; Sankaran, J.; Wohland, T. Fluorescence correlation spectroscopy: The technique and its applications in soft matter. Physical Sciences Reviews 2019, 4, DOI: 10.1515/psr-2017-0104.

(15) Dawes, M. L.; Soeller, C.; Scholpp, S. Studying molecular interactions in the intact organism: fluorescence correlation spectroscopy in the living zebrafish embryo. Histochemistry and Cell Biology 2020, 154, 507–519, DOI: 10.1007/s00418-020-01930-5.

(16) Eggeling, C.; Brand, L.; Ullmann, D.; Jäger, S. Highly sensitive fluorescence detection technology currently available for HTS. Drug Discovery Today 2003, 8, 632–641, DOI: 10.1016/S1359-6446(03)02752-1.

(17) Wachsmuth, M.; Conrad, C.; Bulkescher, J.; Koch, B.; Mahen, R.; Isokane, M.; Pepperkok, R.; Ellenberg, J. High-throughput fluorescence correlation spectroscopy enables analysis of proteome dynamics in living cells. Nature Biotechnology 2015, 33, 384–389, DOI: 10.1038/nbt.3146.

(18) Waithe, D.; Schneider, F.; Chojnacki, J.; Clausen, M. P.; Shrestha, D.; de la Serna, J. B.; Eggeling, C. Optimized processing and analysis of conventional confocal microscopy generated scanning FCS data. Methods 2018, 140-141, 62–73, DOI: 10.1016/j.ymeth.2017.09.010.

(19) Deprey, K.; Becker, L.; Kritzer, J.; Plückthun, A. Trapped! A Critical Evaluation of Methods for Measuring Total Cellular Uptake versus Cytosolic Localization. Bioconjugate Chemistry 2019, 30, 1006–1027, DOI: 10.1021/acs.bioconjchem.9b00112.

(20) Farka, Z.; Mickert, M. J.; Pastucha, M.; Mikušová, Z.; Skládal, P.; Gorris, H. H. Advances in Optical Single-Molecule Detection: En Route to Supersensitive Bioaffinity Assays. Angewandte Chemie International Edition 2020, 59, 10746–10773, DOI: 10.1002/anie.201913924.

(21) Elson, E. L.; Magde, D. Fluorescence correlation spectroscopy. I. Conceptual basis and theory. Biopolymers 1974, 13, 1–27, DOI: 10.1002/bip.1974.360130102.

(22) Magde, D.; Elson, E. L.; Webb, W. W. Fluorescence correlation spectroscopy. II. An experimental realization. Biopolymers 1974, 13, 29–61, DOI: 10.1002/bip.1974.360130103.

(23) Wohland, T.; Maiti, S.; Macháň, R. An Introduction to Fluorescence Correlation Spectroscopy ; 2053-2563; IOP Publishing, 2020; pp 7–1 to 7–23, DOI: 10.1088/978-0-7503-2080-1ch7.

(24) Enderlein, J.; Gregor, I.; Patra, D.; Fitter, J. Art and Artefacts of Fluorescence Correlation Spectroscopy. Current Pharmaceutical Biotechnology 2004, 5, 155–161, DOI: 10.2174/1389201043377020.

(25) Enderlein, J.; Gregor, I.; Patra, D.; Dertinger, T.; Kaupp, U. B. Performance of Fluorescence Correlation Spectroscopy for Measuring Diffusion and Concentration. ChemPhysChem 2005, 6, 2324–2336, DOI: 10.1002/cphc.200500414.

(26) Kim, S. A.; Heinze, K. G.; Schwille, P. Fluorescence correlation spectroscopy in living cells. Nature Methods 2007, 4, 963–973, DOI: 10.1038/nmeth1104.

(27) Petrov, E. P.; Schwille, P. In Standardization and Quality Assurance in Fluorescence Measurements II ; Resch-Genger, U., Ed.; Springer Berlin Heidelberg: Berlin, Heidelberg, 2008; Vol. 6; pp 145–197, DOI: 10.1007/4243_2008_032.

(28) Ries, J.; Petrášek, Z.; García-Sáez, A. J.; Schwille, P. A comprehensive framework for fluorescence cross-correlation spectroscopy. New Journal of Physics 2010, 12, 113009, DOI: 10.1088/1367-2630/12/11/113009.

(29) Macháň, R.; Hof, M. Practical Manual For Fluorescence Microscopy Techniques; PicoQuant GmbH, 2016; p 24.

(30) Bacia, K.; Schwille, P. A dynamic view of cellular processes by in vivo fluorescence auto- and cross-correlation spectroscopy. Methods 2003, 29, 74–85, DOI: 10.1016/S1046-2023(02)00291-8.

(31) Mueller, V.; Honigmann, A.; Ringemann, C.; Medda, R.; Schwarzmann, G.; Eggeling, C. In Methods in Enzymology; Tetin, S. Y., Ed.; Fluorescence Fluctuation Spectroscopy (FFS), Part B; Academic Press, 2013; Vol. 519; pp 1–38, DOI: 10.1016/B978-0-12-405539-1.00001-4.

(32) Altman, R.; Ly, S.; Hilt, S.; Petrlova, J.; Maezawa, I.; Kálai, T.; Hideg, K.; Jin, L.W.; Laurence, T. A.; Voss, J. C. Protective spin-labeled fluorenes maintain amyloid beta peptide in small oligomers and limit transitions in secondary structure. Biochimica et Biophysica Acta (BBA) - Proteins and Proteomics 2015, 1854, 1860–1870, DOI: 10.1016/j.bbapap.2015.09.002.

(33) Khmelinskaia, A.; Mücksch, J.; Conci, F.; Chwastek, G.; Schwille, P. FCS Analysis of Protein Mobility on Lipid Monolayers. Biophysical Journal 2018, 114, 2444–2454, DOI: 10.1016/j.bpj.2018.02.031.

(34) Dunsing, V.; Chiantia, S. A Fluorescence Fluctuation Spectroscopy Assay of Protein-Protein Interactions at Cell-Cell Contacts. Journal of Visualized Experiments: JoVE 2018, DOI: 10.3791/58582.

(35) Qian, H.; Elson, E. L.; Frieden, C. Studies on the structure of actin gels using time correlation spectroscopy of fluorescent beads. Biophysical Journal 1992, 63, 1000–1010, DOI: 10.1016/S0006-3495(92)81686-7.

(36) Günther, J.-P.; Börsch, M.; Fischer, P. Diffusion Measurements of Swimming Enzymes with Fluorescence Correlation Spectroscopy. Accounts of Chemical Research 2018, 51, 1911–1920, DOI: 10.1021/acs.accounts.8b00276.

(37) Milon, S.; Hovius, R.; Vogel, H.; Wohland, T. Factors influencing fluorescence correlation spectroscopy measurements on membranes: simulations and experiments. Chemical Physics 2003, 288, 171–186, DOI: 10.1016/S0301-0104(03)00018-1.

(38) Ries, J.; Schwille, P. New concepts for fluorescence correlation spectroscopy on membranes. Physical Chemistry Chemical Physics 2008, 10, 3487–3497, DOI: 10.1039/B718132A.

(39) Baum, M.; Erdel, F.; Wachsmuth, M.; Rippe, K. Retrieving the intracellular topology from multi-scale protein mobility mapping in living cells. Nature Communications 2014, 5, 4494, DOI: 10.1038/ncomms5494.

(40) Dunsing, V. Fluorescence fluctuation spectroscopy techniques to quantify molecular interactions and dynamics in complex biological systems. Ph.D. thesis, Universität Potsdam, Potsdam, Germany, 2020; 10.25932/publishup-47849.

(41) Lahtinen, S.; Krause, S.; Arppe, R.; Soukka, T.; Vosch, T. Upconversion Cross-Correlation Spectroscopy of a Sandwich Immunoassay. Chemistry (Weinheim an Der Bergstrasse, Germany) 2018, 24, 9229–9233, DOI: 10.1002/chem.201801962.

(42) Murray, R. A.; Qiu, Y.; Chiodo, F.; Marradi, M.; Penadés, S.; Moya, S. E. A Quantitative Study of the Intracellular Dynamics of Fluorescently Labelled Glyco-Gold Nanoparticles via Fluorescence Correlation Spectroscopy. Small 2014, 10, 2602–2610, DOI: 10.1002/smll.201303604.

(43) Garai, K.; Sahoo, B.; Kaushalya, S. K.; Desai, R.; Maiti, S. Zinc Lowers Amyloidβ Toxicity by Selectively Precipitating Aggregation Intermediates. Biochemistry 2007, 46, 10655–10663, DOI: 10.1021/bi700798b.

(44) Ferrand, P.; Wenger, J.; Rigneault, H. In Single Molecule Analysis, Methods and Protocols; Peterman, E. J. G., Wuite, G. J. L., Eds.; Methods in Molecular Biology; Humana Press, Springer, 2011; pp 181–195, DOI: 10.1007/978-1-61779-282-3_10.

(45) Carl Zeiss MicroImaging GmbH, LSM 710, LSM 780, LSM 710 NLO, LSM 780 NLO and ConfoCor 3 Operating Manual, Chapter 9: Confocor 3 (FCS). 2010.

(46) Widengren, J.; Rigler, R. Fluorescence correlation spectroscopy as a tool to investigate chemical reactions in solutions and on cell surfaces. Cellular and Molecular Biology (Noisy-Le-Grand, France) 1998, 44, 857–879.

(47) Saffarian, S.; Elson, E. L. Statistical Analysis of Fluorescence Correlation Spectroscopy: The Standard Deviation and Bias. Biophysical Journal 2003, 84, 2030–2042, DOI: 10.1016/S0006-3495(03)75011-5.

(48) Wohland, T.; Rigler, R.; Vogel, H. The Standard Deviation in Fluorescence Correlation Spectroscopy. Biophysical Journal 2001, 80, 2987–2999, DOI: 10.1016/S0006-3495(01)76264-9.

(49) Persson, G.; Thyberg, P.; Sandén, T.; Widengren, J. Modulation Filtering Enables Removal of Spikes in Fluorescence Correlation Spectroscopy Measurements without Affecting the Temporal Information. The Journal of Physical Chemistry B 2009, 113, 8752–8757, DOI: 10.1021/jp902538b.

(50) Ries, J.; Bayer, M.; Csućs, G.; Dirkx, R.; Solimena, M.; Ewers, H.; Schwille, P. Automated suppression of sample-related artifacts in Fluorescence Correlation Spectroscopy. Optics Express 2010, 18, 11073–11082, DOI: 10.1364/OE.18.011073.

(51) Miller, A. E.; Hollars, C. W.; Lane, S. M.; Laurence, T. A. Fluorescence Cross-Correlation Spectroscopy as a Universal Method for Protein Detection with Low False Positives. Analytical Chemistry 2009, 81, 5614–5622, DOI: 10.1021/ac9001645.

(52) Wood, C.; Huff, J.; Marshall, W.; Yu, E. Q.; Unruh, J.; Slaughter, B.; Wiegraebe, W. Fluorescence correlation spectroscopy as tool for high-content-screening in yeast (HCS-FCS). Single Molecule Spectroscopy and Imaging IV. 2011; pp 54–68, DOI: 10.1117/12.873947.

(53) Moen, E.; Bannon, D.; Kudo, T.; Graf, W.; Covert, M.; Van Valen, D. Deep learning for cellular image analysis. Nature Methods 2019, 16, 1233–1246, DOI: 10.1038/s41592-019-0403-1.

(54) Ren, S. S. Imaging Fluorescence Correlation Spectroscopy Analysis Using Convolutional Neural Networks. Thesis, 2022.

(55) Wohland, T.; Tang, W. H.; Sim, S. R.; Aik, D.; Röllin, A. Deep learning approaches for imaging fluorescence correlation spectroscopy parameter estimation with limited data sets. Biophysical Journal 2022, 121, 533a, DOI: 10.1016/j.bpj.2021.11.2808.

(56) Böhmer, M.; Wahl, M.; Rahn, H.-J.; Erdmann, R.; Enderlein, J. Time-resolved fluorescence correlation spectroscopy. Chemical Physics Letters 2002, 353, 439–445, DOI: 10.1016/S0009-2614(02)00044-1.

(57) Macháň, R.; Kapusta, P.; Hof, M. Statistical filtering in fluorescence microscopy and fluorescence correlation spectroscopy. Analytical and Bioanalytical Chemistry 2014, 406, 4797–4813, DOI: 10.1007/s00216-014-7892-7.

(58) Ronneberger, O.; Fischer, P.; Brox, T. In Medical Image Computing and Computer-Assisted Intervention –MICCAI 2015 ; Navab, N., Hornegger, J., Wells, W. M., Frangi, A. F., Eds.; Lecture Notes in Computer Science; Springer International Publishing: Cham, 2015; Vol. 9351; pp 234–241, DOI: 10.1007/978-3-319-24574-4_28.

(59) Falk, T. et al.. U-Net: deep learning for cell counting, detection, and morphometry. Nature Methods 2019, 16, 67–70, DOI: 10.1038/s41592-018-0261-2.

(60) von Chamier, L. et al.. Democratising deep learning for microscopy with ZeroCostDL4Mic. Nature Communications 2021, 12, 2276, DOI: 10.1038/s41467-021-22518-0.

(61) Maddalena, L.; Antonelli, L.; Albu, A.; Hada, A.; Guarracino, M. R. Artificial Intelligence for Cell Segmentation, Event Detection, and Tracking for Label-Free Microscopy Imaging. Algorithms 2022, 15, 313, DOI: 10.3390/a15090313.

(62) Punn, N. S.; Agarwal, S. Modality specific U-Net variants for biomedical image segmentation: a survey. Artificial Intelligence Review 2022, 55, 5845–5889, DOI: 10.1007/s10462-022-10152-1.

(63) Jimenez-Perez, G.; Alcaine, A.; Camara, O. U-Net Architecture for the Automatic Detection and Delineation of the Electrocardiogram. 2019 Computing in Cardiology (CinC). 2019; pp Page 1–Page 4, DOI: 10.22489/CinC.2019.284, ISSN: 2325-887X.

(64) Oh, S. L.; Ng, E. Y. K.; Tan, R. S.; Acharya, U. R. Automated beat-wise arrhythmia diagnosis using modified Unet on extended electrocardiographic recordings with heterogeneous arrhythmia types. Computers in Biology and Medicine 2019, 105, 92–101, DOI: 10.1016/j.compbiomed.2018.12.012.

(65) Cheng, X.; Liu, D.; Lu, J.; Wei, L.; Hu, A.; Lei, J.; Zou, Z.; Zou, X.; Jiang, Q. Efficient hardware design of a deep U-net model for pixel-level ECG classification in healthcare device. Microelectronics Journal 2022, 126, 105492, DOI: 10.1016/j.mejo.2022.105492.

(66) Dmitrieva, M.; Lefebvre, J.; delas Peñas, K.; Zenner, H. L.; Richens, J.; St Johnston, D.; Rittscher, J. Short Trajectory Segmentation with 1D UNET Framework: Application to Secretory Vesicle Dynamics. 2020 IEEE 17th International Symposium on Biomedical Imaging (ISBI). 2020; pp 891–894, DOI: 10.1109/ISBI45749.2020.9098426,ISSN: 1945-8452.

(67) Guo, S.; Mayerhöfer, T.; Pahlow, S.; Hübner, U.; Popp, J.; Bocklitz, T. Deep learning for ‘artefact’ removal in infrared spectroscopy. Analyst 2020, 145, 5213–5220, DOI: 10.1039/D0AN00917B.

(68) Gebrekidan, M. T.; Knipfer, C.; Braeuer, A. S. Refinement of spectra using a deep neural network: Fully automated removal of noise and background. Journal of Raman Spectroscopy 2021, 52, 723–736, DOI: 10.1002/jrs.6053.

(69) Alexa Fluor™ 488 NHS Ester (Succinimidyl Ester). https://www.thermofisher.com/order/catalog/product/de/en/A20000.

(70) Moreno-Torres, J. G.; Raeder, T.; Alaiz-Rodríguez, R.; Chawla, N. V.; Herrera, F. A unifying view on dataset shift in classification. Pattern Recognition 2012, 45, 521–530, DOI: 10.1016/j.patcog.2011.06.019.

(71) Pignataro, M. F.; Herrera, M. G.; Dodero, V. I. Evaluation of Peptide/Protein Self-Assembly and Aggregation by Spectroscopic Methods. Molecules 2020, 25, 4854, DOI: 10.3390/molecules25204854.

(72) Kitamura, A.; Kinjo, M. State-of-the-Art Fluorescence Fluctuation-Based Spectroscopic Techniques for the Study of Protein Aggregation. International Journal of Molecular Sciences 2018, 19, 964, DOI: 10.3390/ijms19040964.

