## Supplementary Note, Supplementary Tables S1-S3, Supplementary Figures S1-S4 for "Neural Network Informed Photon Filtering Reduces Artifacts in Fluorescence Correlation Spectroscopy Data"

*Simulation experiments.* We performed Monte Carlo simulations of FCS fluorescence time-series based on the NANOSIMPY v30/03/2020 module[1] in PYTHON v3.9.5.[2] For fluorescence time-series without artifacts, we simulated 16,384 time steps (step size: 1 ms) of the 2-dimensional stochastic Brownian motion of 500–3.500 freely moving particles in a box of 3.000 nm × 3.000 nm and a central observation spot with a full width at half-maximum of 250 nm, as described previously.[1,3,4] We simulated 300 time-series per 10 groups of diffusion coefficients determining the time-series segments without artifacts $D_{\mathrm{mol}} = \{0.069, 0.08, 0.1, 0.2, 0.4, 0.6, 1.0, 3.0, 10, 50\}\,\mu\mathrm{m}^2\,\mathrm{s}^{-1}$ (corresponding to $\tau_{\mathrm{mol}} = \{163, 141, 113, 56.4, 28.2, 18.8, 11.3, 3.76, 1.13, 0.23\}\,\mathrm{ms}$) We used the same method to simulate three groups of clusters determining time-series segments with peak artifacts (3 particles and $D_{\mathrm{clust}} = 1\,\mu\mathrm{m}^2\,\mathrm{s}^{-1}$, 7 particles and $D_{\mathrm{clust}} = 0.1\,\mu\mathrm{m}^2\,\mathrm{s}^{-1}$, 10 particles and $D_{\mathrm{clust}} = 0.01\,\mu\mathrm{m}^2\,\mathrm{s}^{-1}$). To create fluorescence time-series with peak artifacts, we scaled the clusters by a factor of 5.000–10.000 and added one to each time-series without artifacts, yielding 30 groups for model training. To create a segmentation vector showing the location of the artifacts, we applied a cluster threshold of 0.04 to the non-scaled cluster time-series (based on preliminary plots similar to Figure 2B). We simulated additional 6000 time-series for training and validation of the neural network. We employed 100 previously unseen time-series from all 30 groups as test data for the neural network and used a subset of 9 groups to test the correction methods (see Figure 2).

*Applied experiments.* The first peak artifact measurements were produced by mixing 20 nM of the small dye Alexa Fluor 488 (*Thermo Fisher Scientific*) with 10 µM of slow, large (100 nm) unilamellar vesicles (LUVs) labelled with DiO in the membrane (0.1 % mol lipid:dye). The corresponding control measurements were 20 nM Alexa Fluor 488 in solution. We prepared the LUVs by the extrusion method: 1−palmitoyl−2−oleoyl−glycero−3−phosphocholine (POPC, *Avanti Polar Lipids*) was vacuum dried on a glass vial for 1 h and resuspended on phosphate buffered saline to form lipid vesicles. The vesicles were extruded 30 times through 100 nm pore size polycarbonate membranes using a *Mini-Extruder* (*Avanti Polar Lipids*). The second peak artifacts measurements were produced by 20 nM Trypanosoma brucei-PEX5 N-term fused to eGFP in solution, and the corresponding control measurements by 5 nM Homo sapiens-PEX5 N-term fused to eGFP in solution. The detailed sample preparation is described elsewhere.[5,6]

We prepared the samples on #1.5 coverslips mounted on *Attofluor Cell Chambers* (*Thermo Fisher Scientific*). We acquired the data on a *MicroTime 200* microscope (*PicoQuant*) equipped with an *Olympus UPlanSApo* 60× 1.2*NA* water immersion objective lens and a *HydraHarp 400* time-correlated single photon counting (TCSPC) module (*PicoQuant*). Excitation was achieved with a 488 nm pulsed laser (*PicoQuant*) with a power of 5 mW measured at the sample plane. We used a 20 nM solution of Alexa Fluor for calibrating the correction collar. One TCSPC measurement had a length of 10 s for AF488 experiments and 20 s for PEX5 experiments. In total, the dataset sizes were $n_{\text{AF488}} = 424$, $n_{\text{AF488+DiOLUVs}} = 440$, $n_{\text{Hs-PEX5-eGFP}} = 250$, and $n_{\text{Tb-PEX5-eGFP}} = 250$.

*Supervised machine learning.* For training, we used simulated fluorescence time-series with peak artifacts as features, and simulated segmentation vectors (0 = no artifact, 1 = peak artifact). The total dataset was split into 3 parts: $4,800$ time-series for training, $1,200$ time-series for validation, and $3,000$ time-series for testing. We implemented a 1-dimensional U-Net[7,8] in TensorFlow 2 v2.5.0[9,10] including pre-processing functions from Scikit-learn v0.24.2.[11] All model training runs were scheduled and tracked, including relevant metadata, using mlflow v1.19.0.[12,13] For hyperparameter tuning, we used mlflow v1.19.0 and a random search algorithm via the TensorBoard HParams API v2.5.0. To support the reproducibility of the machine learning method of this study, we included the machine learning summary table (Table S1) in the supporting information as per DOME recommendations.[14] All models with a validation precision at a 0.5 prediction threshold $prec_{0.5} > 0.85$ and a validation sensitivity at a 0.5 prediction threshold $sens_{0.5} > 0.85$ were further evaluated as predictors in the automated correction pipeline. We avoided manual annotation of the application data by comparing fit outcome distributions instead of prediction performance. We also compared the U-Net predictor against a baseline predictor consisting of thresholding after applying SKLEARN.PREPROCESSING.ROBUSTSCALER. For Figures 4C and 5C, we chose the best of three thresholds, which were 1.5, 2 and 2.5 for AF488 data and 5, 7 and 10 for PEX5 data (see Figure S4). The model version `0cd20` (large model, 200 MB) performed best in simulations and applied experiments (see Figures 4B-C and 5B-C). The smaller model version `ff67b` (small model, 14 MB) might be preferable if the model size is crucial, as in mobile deployment (for comparison see Table S2 and Figure S4).

*Correction pipeline and FCS Analysis.* We wrote two PYTHON v3.9.5 implementations for the correction methods based on NUMPY v1.21.5[15] and PANDAS v1.3.5.[16,17] For simulations, we applied *cut and stitch* and *set to zero* corrections on the simulated time-series directly. We autocorrelated the simulated time-series using the Python package MULTIPLETAU v0.3.3.[18–20] For the TCSPC data from the applied experiments, we used a fork of the TTTR2XFCS algorithm[21] as implemented in FOCUS-POINT v1.16.203[22,23] for autocorrelation. We added custom methods to extract binned time-series, predict peak artifacts on these extracted time-series using the trained neural network, and map the segmentation back to the TCSPC arrival times to perform the *cut and stitch* and *set to zero* correction methods.

We fitted the data using FOCUS-POINT v1.16.203. We evaluated both 1 and 2 species fits for all simulations and applied experiments using PANDAS v1.3.5 for statistics such as median and interquartile range (IQR) and SEABORN v0.11.2 for fit outcome distribution plots. For Figures 2B, 4B-C and 5B-C, we only displayed the distributions closest to the expected fit outcome value, all fit outcome distributions are shown in Figures S3 and S4. For a list of FCS fit equations and parameters see Table S3. All transit time distributions are plotted on a log scale to capture the log Gaussian nature, which leads to higher variance for higher transit time values.[1]

### Supplementary Tables

Table S1: Report on supervised machine learning according to the DOME recommendations[14]

| **DOME** | Version | 1.0 |
|---|---|---|
| **Data: *Nanosimpy simulations*** | Provenance | Own simulations. Total of $N = 147,456,000$ time steps segmented in peak artifact vs no artifact (9000 time-series with $16,384$ time steps each). $N_{pos} = 22,688,854$, $N_{neg} = 124,767,146$. Not previously used. |
| | Dataset splits | $4,800$ time-series for training ($N_{train,tot} = 78,643,200$, $N_{train,pos} = 11,969,767$, $N_{train,neg} = 66,673,433$, 15.22% positives on train set), $1,200$ time-series for validation ($N_{val,tot} = 19,660,800$, $N_{val,pos} = 3,162,854$, $N_{val,neg} = 16,497,946$, 16.09% positives on validation set), $3,000$ time-series for testing ($N_{test,tot} = 49,152,000$, $N_{test,pos} = 7,556,233$, $N_{test,neg} = 41,595,767$, 15.37% positives on test set) |
| | Redundancy between data splits | Independent simulations. Train and test used the same list of 10 types of $D_{\mathrm{mol}}$ and 3 types of $D_{\mathrm{clust}}$, but different numbers of particles / molecules. Validation used the same list of $D_{\mathrm{mol}}$, but only a subset of $D_{\mathrm{clust}}$. |
| | Availability of data | Code: `ttps://doi.org/10.5281/zenodo.8137220`. Data: `https://zenodo.org/record/8074408`. Literate simulation code: `https://aseltmann.github.io/fluotracify/data/LabBook-all.html#sec-2-2` |

Continuation of Table S1

| | | |
|---|---|---|
| **Optimization** | Algorithm | Fully convolutional neural network, specifically 1-D U-NET[7,8], own implementation with TensorFlow 2 v2.5.0. Optimizer: Adam. Loss: binary cross entropy (loss for each time step) + dice loss (loss for each time-series). See Figure 3 for overview of U-Net architecture. |
| | Meta-predictions | No. |
| | Data encoding | Replace NaN values with zero. Scale time-series (see Table S2). Pad the end of the time-series with the median up to a length of the next biggest power of 2 (Only training: Crop length to $14,000$, then pad to $16,384$). |
| | Parameters | $p_{\texttt{0cd20}} = 17,030,783$, $p_{\texttt{ff67b}} = 844,123$. We used Mlflow v1.19.0 for hyperparameter selection and logging. MLflow performed 2 runs for each random combination of a set of hyperparameters for 20 epochs, and averaged the performance metrics. We chose all hyperparameter combinations with both a validation precision $> 0.85$ and a validation recall $> 0.85$ (10 models) and performed a dedicated training from scratch for 100 epochs per model. We selected 2 models based on their performance in the whole pipeline (see Table S2). |
| | Features | The model accepts arbitrary powers of 2 as input length for time-series ($f = 1,024, 2,048, \ldots$). By including the padding step as described above, intensity time-series of arbitrary lengths can be segmented. Input data during training: $f = 16,384$. No feature selection. |

Continuation of Table S1

| | | |
|---|---|---|
| | Fitting | We observed no overfitting during hyperparameter search when comparing loss and validation loss over epochs. Due to a misconfiguration in mlflow, no training metrics over epochs were saved for the re-training of the 10 best models. Since no large gap was present between loss and validation loss after the 100th epoch according to training logs, and the models performed well when applied in the correction pipeline for biological experiments, we assume no large influence of overfitting. |
| | Regularization | No. |
| | Availibility of configuration | Yes. Training code: `https://doi.org/10.5281/zenodo.8137220`. Literate hyperparameter training code: `https://aseltmann.github.io/fluotracify/data/LabBook-all.html#sec-2-4`. Literate final model training code: `https://aseltmann.github.io/fluotracify/data/LabBook-all.html#sec-2-6`. Free use license: Apache License 2.0. |
| **Model** | Interpretability | Black box, as correlation between input and output is masked. No attempt was made to make the model transparent. |
| | Output | Segmentation, i. e. probability of each time step to be artifactual / dominated by a peak artifact in FCS measurements. |
| | Execution time | HPC inference time (one 16,384-length time-series): version `0cd20`: 22 ms, version `ff67b`: 15 ms. |
| | Availibility of software | Yes. Models: `https://doi.org/10.5281/zenodo.8137129`. Loadable via MLFLOW (`https://www.mlflow.org`). |

Continuation of Table S1

| | | |
|---|---|---|
| **Evaluation** | Evaluation method | Independent test dataset. Additionally biophysical control in combination with *cut and stitch* correction of fluorescence time-series and successful restoration of fitted parameters $(\tau, N)$ via FCS analysis in independent biological experiments. |
| | Performance measures | ROC-AUC score. With a prediction threshold of 0.5: F1 score, precision, recall. |
| | Comparison | Manual thresholding after robust scaling as control. |
| | Confidence | Not calculated. |
| | Availability of evaluation | Yes. Code and plots: `https://doi.org/10.5281/zenodo.8137220`. Literate final model evaluation code: `https://aseltmann.github.io/fluotracify/data/LabBook-all.html#sec-2-6`. Free use license: Apache License 2.0 for software, Creative Commons Attribution 4.0 International License for non-software. |

Table S2: Machine learning model hyperparameters and test performance

| | **Model version `0cd20`** | **Model version `ff67b`** |
|---|---|---|
| Parameters | $17,030,783$ | $844,123$ |
| Model size | 200 MB | 14 MB |
| Training batch size | 15 | 28 |
| U-Net levels | 6 | 5 |
| Filters per level | down: $[23, 46, 92, 184, 368, 512]$, center: 512, up: $[512, 368, 184, 92, 46, 23]$ | down: $[6, 12, 24, 48, 96]$, center: 192, up: $[96, 48, 24, 12, 6]$ |
| Input size | 16384 | 16384 |
| Learning rate | polynomial decay (7th power), start = 0.0305 | linear decay, start = 0.0553 |
| `MaxPooling1D` size | 4 | 4 |
| Scaler | quantile transformation (gaussian pdf) | min-max |
| Train loss after last epoch | 0.1275 | 0.0890 |
| Validation loss after last epoch | 0.1678 | 0.1286 |
| Test AUC | 0.9614 | 0.9756 |
| Test F1 (threshold 0.5) | 0.8863 | 0.9201 |
| Test precision (thr 0.5) | 0.9070 | 0.9301 |
| Test recall (thr 0.5) | 0.8665 | 0.9104 |
| Avg HPC inference time (one $16,384$-length time-series, $n = 3000$) | 22 ms | 15 ms |

Table S3: FCS fitting model equations and parameters

| Data | Model equation | Parameters |
| --- | --- | --- |
| overall equation: $G_N(\tau) = O_f + G_N(0)\left[G_D(\tau) \cdot G_T(\tau)\right]$ with offset $O_f$, correlation function amplitude $G_N(0)$, translational diffusing component $G_D(\tau)$ ($G_{2D}(\tau)$ for 2D and $G_{3D}(\tau)$ for 3D) and triple state $G_T(\tau)$ | | |
| Nanosimpy simulations *without artifacts* | 2D equation, no triplet, 1 species: $G_{2D}(\tau) = \left(1 + \left(\frac{\tau}{\tau_{D_k}}\right)^{\alpha}\right)^{-1}$ | <u>fixed parameters</u>: anomaly parameter $\alpha = 1$, $x_{\min} = 1$, $x_{\max} = 8192$ <u>floating parameters</u>: $O_f$, $G_N(0)$, lateral transit time $\tau_{D_k}$ |
| Nanosimpy simulations *with peak artifacts* | 2D equation, no triplet, 2 separate fits for 1 and 2 species: $G_{2D}(\tau) = \sum_{k=1}^{D_s} A_k \left(1 + \left(\frac{\tau}{\tau_{D_k}}\right)^{\alpha^k}\right)^{-1}$ | <u>fixed parameters</u>: $\alpha = 1$, diffusion species $D_s = \{1, 2\}$, $x_{\min} = 1$, only *averaging* correction: $x_{\max} = 1024$, other processing: $x_{\max} = 8192$ <u>floating parameters</u>: $O_f$, $G_N(0)$, $\tau_{D_k}$, relative species fraction $A_k$ (for $D_s = 1$, $A_1 = 1$, else $\sum D_s A_k = 1$) |
| AF488 data | 3D equation, no triplet, 2 separate fits for 1 and 2 species: $G_{3D}(\tau) = \sum_{k=1}^{D_s} A_k \left(1 + \left(\frac{\tau}{\tau_{D_k}}\right)^{\alpha^k}\right)^{-1} \cdot \left(1 + \left(\frac{\tau}{AR_k^2 \tau_{D_k}}\right)\right)^{-1/2}$ | <u>fixed parameters</u>: $\alpha = 1$, $D_s = \{1, 2\}$, aspect ratio factor $AR_k = 5$, $x_{\min} = 0.001$, only *averaging* correction: $x_{\max} = 0.5$, other processing, time-series without peak artifacts: $x_{\max} = 100$, other processing, time-series with peak artifacts: $x_{\max} = 500$ <u>floating parameters</u>: $O_f$, $G_N(0)$, $\tau_{D_k}$, $A_k$ |

Continuation of Table S3

| | | |
|---|---|---|
| PEX5 data | 3D equation, 1 triplet state, 2 separate fits for 1 and 2 species: $G_{3D}(\tau) = \sum_{k=1}^{D_s} A_k \Big(1 + \big(\frac{\tau}{\tau_{D_k}}\big)^{\alpha^k}\Big)^{-1} \cdot \Big(1 + \big(\frac{\tau}{AR_k^2 \tau_{D_k}}\big)\Big)^{-1/2}$ <br> $G_T(\tau) = 1 - T + T \cdot \exp\big(-\frac{\tau}{\tau_T}\big)$ | fixed parameters: $\alpha = 1$, $D_s = \{1, 2\}$, $AR_k = 6$, triplet decay time $\tau_T = 0.04$ (as used in Galiani et al.[5]), $x_{\min} = 0.001$, $x_{\max} = 1000$ <br> floating parameters: $O_f$, $G_N(0)$, $\tau_{D_k}$, $A_k$, triplet fraction size $T$ |

### Supplementary Figures

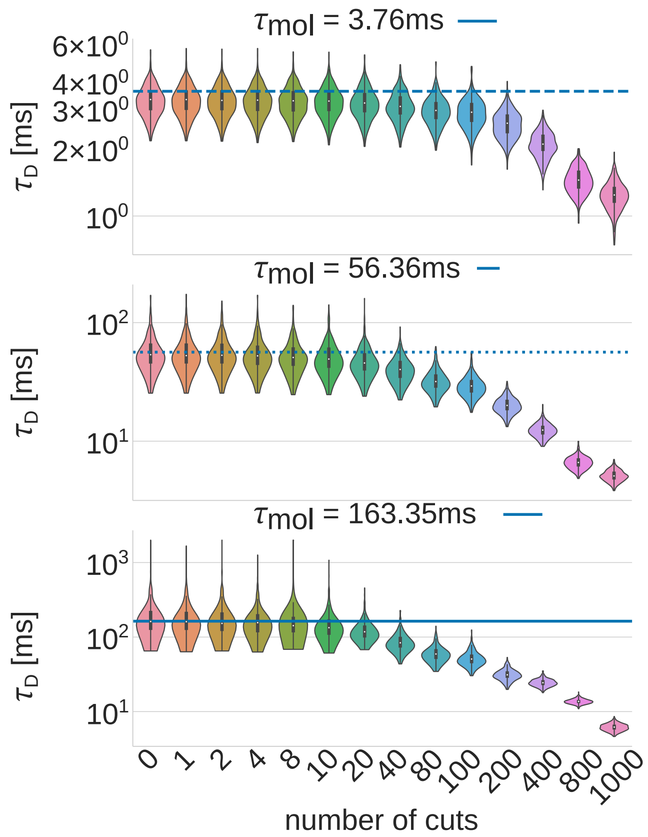

Figure S1: Effect of an increasing number of randomly placed cuts on simulated fluorescence time-series *without artifacts* (notation: three simulated molecule transit times $\tau_{\text{mol}}$ determine the shape of the simulated fluorescence time-series, and the fitted transit times $\tau_D$ result from correlation and a one species FCS fit. Further parameters: $n = 300$, time-series length $=$ 16 384 ms, step size $=$ 1 ms). After cutting, we randomly shuffled the parts and stitched them together. The $\tau_D$ distribution shifts after 40 cuts for $\tau_{\text{mol}} = 3.76$ ms, 20 cuts for $\tau_{\text{mol}} = 56.46$ ms, and 10 cuts for $\tau_{\text{mol}} = 163.35$ ms. This illustrates more prominent 'stitching artifacts' for higher $\tau_{\text{mol}}$ (and thus for lower diffusion coefficients $D_{\text{mol}}$). Furthermore, more cuts should be possible for longer time-series and higher temporal resolution (smaller step size).

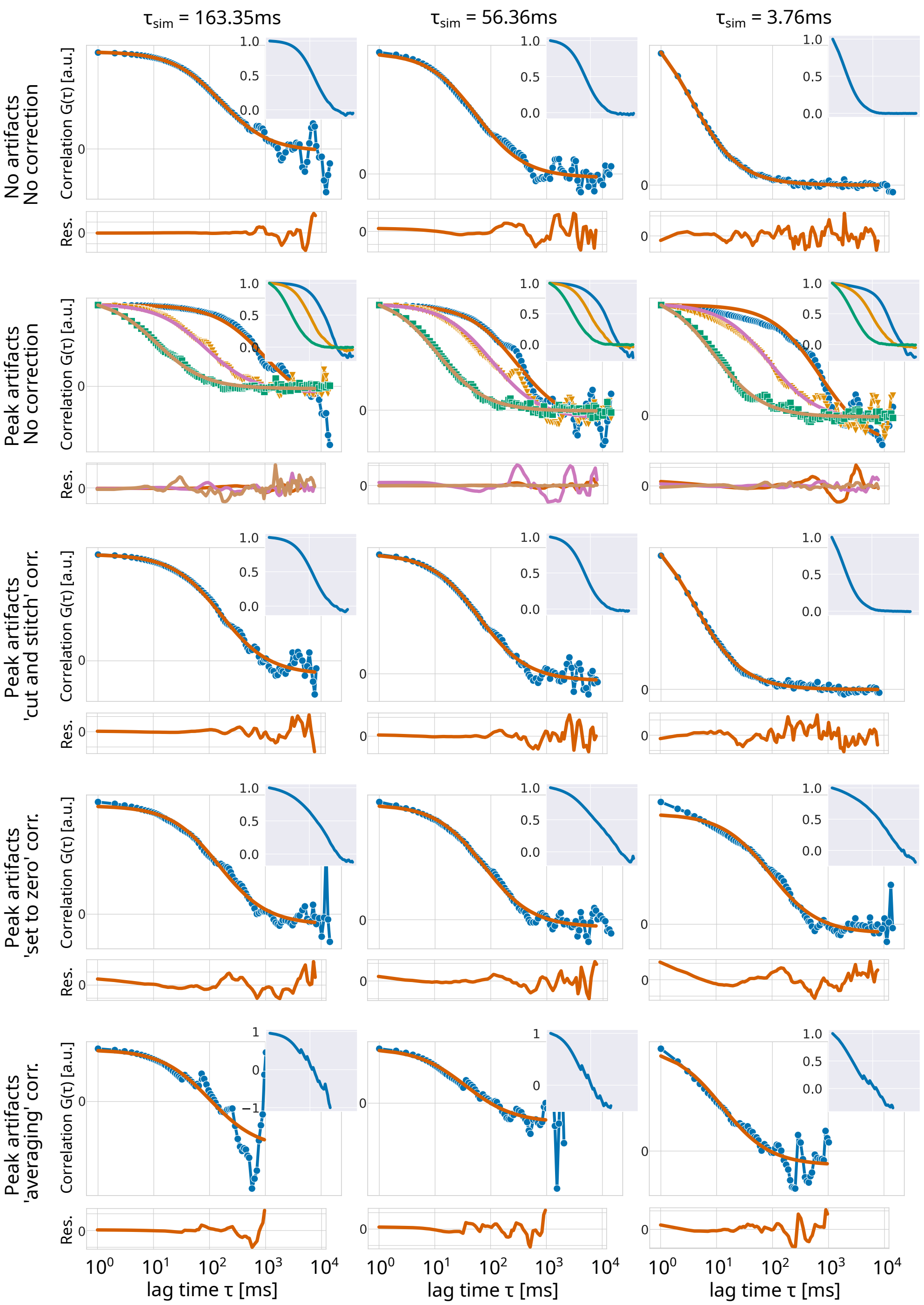

Figure S2: The graphs display how the correction methods (labelled on the left) impact three sets of simulated time-series with varying simulated transit times $\tau_{\mathrm{mol}}$ (labelled on top). We used the simulated segmentation vectors to identify the artifacts to make this comparison independent of the prediction methods. For each category, the upper plot shows one illustrative correlation (marked line) and the corresponding 1-component fit (solid line), the lower plot shows the corresponding residuals, and the inset shows the average, max-abs normalized correlation of all 300 simulated time-series in this group. For the category 'Peak artifacts, No correction', three examples have been chosen to represent the three cluster speeds which dominate the respective correlations (11 ms in green, 112 ms in orange, 1127 ms in blue) and the inset showing the average of all 100 simulated time-series in each subgroup. Note: Because the simulated number of molecules differed for each time-series, only the lag times are comparable between categories. For clarity, we removed the tick labels on the ordinate except for $G = 0$. For a quantitative comparison of fit outcomes see Figure 2.

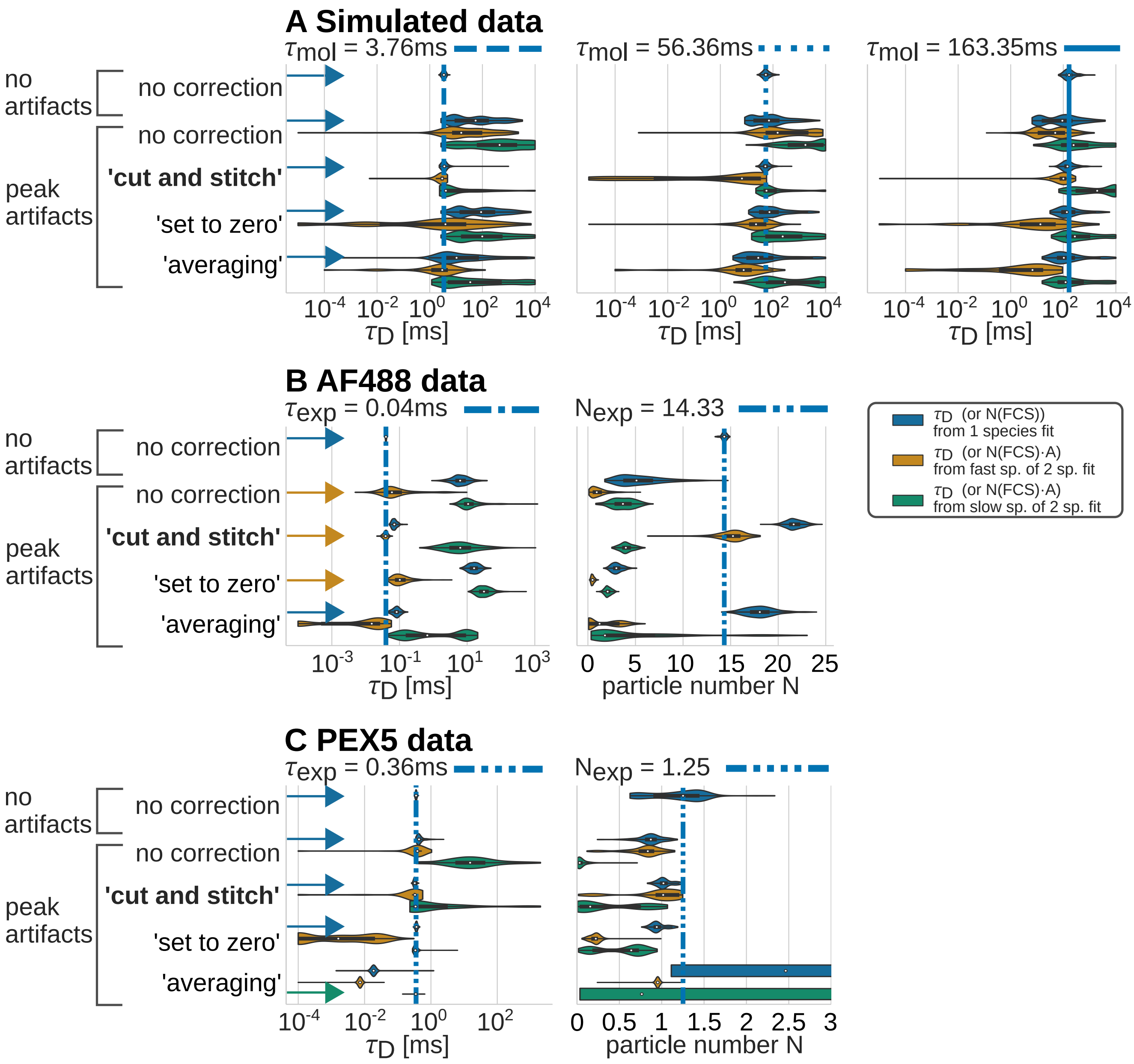

Figure S3: Extended fit outcomes for correction method comparisons of Figures 2B, 4B, and 5B. We displayed transit time $\tau_D$ and particle number $N$ distributions after FCS correlation and fit. All groups with peak artifacts yielded three distributions (1 species fit (blue), fast sp. of 2 sp. fit (orange), slow sp. of 2 sp. fit (green)). Table S3 holds the corresponding FCS fit equations. For the Main Figures, we chose the most charitable fit outcomes with the closest fit between expected values and outcome distribution, as denoted by the arrows next to the labels on the left side. **A** Simulation experiments. Expected transit times $\tau_{\text{mol}}$ from simulations, segmentation from simulations, $n = 300$. **B-C** Applied experiments. Expected transit times $\tau_{\text{exp}}$ and particle numbers $N_{\text{exp}}$ from the group 'No artifacts, no correction' (median of 1 component fits), segmentation from U-Net predictor, $n_{\text{AF488}} = 424$ (no artifacts), $n_{\text{AF488+DiOLUVs}} = 440$ (peak artifacts), $n_{\text{Hs-PEX5-eGFP}} = 250$ (no artifacts), and $n_{\text{Tb-PEX5-eGFP}} = 250$ (peak artifacts).

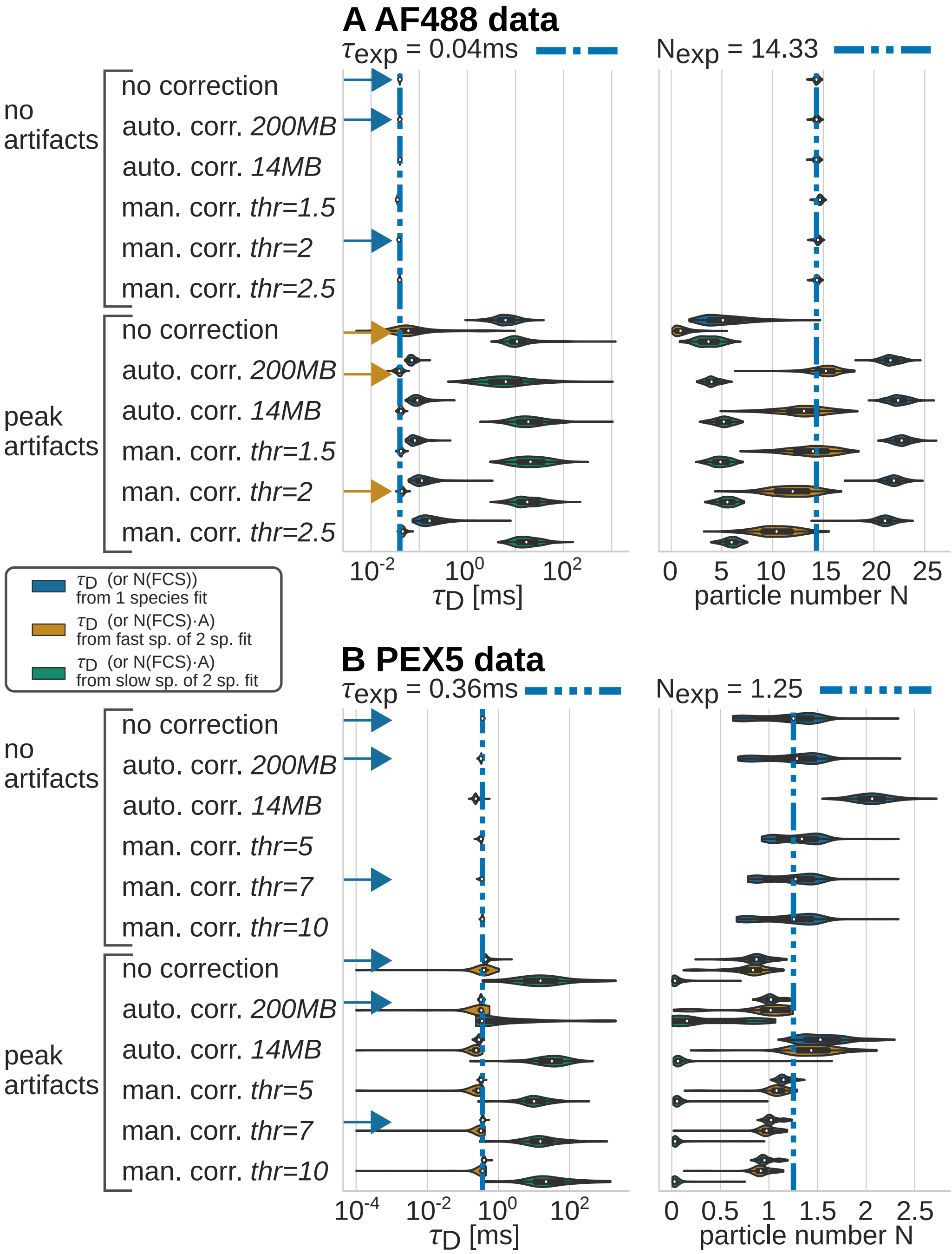

Figure S4: Extended fit outcomes for prediction method comparisons of Figures 4C and 5C for **A** AF488 data and **B** PEX5 data. We displayed transit time $\tau_D$ and particle number $N$ distributions after FCS correlation and fit. We compared five correction pipelines featuring two automated prediction methods (U-Net version `0cd20` (200 MB) and `ff67b` (14 MB)) and three manual prediction methods. For the manual ones, we applied robust scaling and chose three well-fitting thresholds by trial and error. All pipelines used the *cut and stitch* correction method. All groups with peak artifacts yielded three distributions (1 species fit (blue), fast sp. of 2 sp. fit (orange), slow sp. of 2 sp. fit (green)). Table S3 holds the corresponding FCS fit equations. For the Main Figures, we chose one automated and one manual correction pipeline with the most charitable fit outcomes, as denoted by the arrows next to the labels on the left side.

### References

(1) Waithe, D.; Schneider, F.; Chojnacki, J.; Clausen, M. P.; Shrestha, D.; de la Serna, J. B.; Eggeling, C. Optimized processing and analysis of conventional confocal microscopy generated scanning FCS data. *Methods* **2018**, *140-141*, 62–73.

(2) Rossum, G. v.; Drake, F. L. *The Python language reference*, release 3.0.1 [repr.] ed.; Python documentation manual / Guido van Rossum; Fred L. Drake [ed.] Pt. 2; Python Software Foundation: Hampton, NH, 2010.

(3) Pinkwart, K.; Schneider, F.; Lukoseviciute, M.; Sauka-Spengler, T.; Lyman, E.; Eggeling, C.; Sezgin, E. Nanoscale dynamics of cholesterol in the cell membrane. *Journal of Biological Chemistry* **2019**, *294*, 12599–12609.

(4) Schneider, F.; Waithe, D.; Clausen, M. P.; Galiani, S.; Koller, T.; Ozhan, G.; Eggeling, C.; Sezgin, E. Diffusion of lipids and GPI-anchored proteins in actin-free plasma membrane vesicles measured by STED-FCS. *Molecular Biology of the Cell* **2017**, *28*, 1507–1518.

(5) Galiani, S.; Reglinski, K.; Carravilla, P.; Barbotin, A.; Urbančič, I.; Ott, J.; Sehr, J.; Sezgin, E.; Schneider, F.; Waithe, D.; Hublitz, P.; Schliebs, W.; Erdmann, R.; Eggeling, C. Diffusion and interaction dynamics of the cytosolic peroxisomal import receptor PEX5. *Biophysical Reports* **2022**, *2*, 100055.

(6) Schliebs, W.; Saidowsky, J.; Agianian, B.; Dodt, G.; Herberg, F. W.; Kunau, W.-H. Recombinant Human Peroxisomal Targeting Signal Receptor PEX5: STRUCTURAL BASIS FOR INTERACTION OF PEX5 WITH PEX14*. *Journal of Biological Chemistry* **1999**, *274*, 5666–5673.

(7) Ronneberger, O.; Fischer, P.; Brox, T. In *Medical Image Computing and Computer-Assisted Intervention – MICCAI 2015*; Navab, N., Hornegger, J., Wells, W. M., Frangi, A. F., Eds.; Lecture Notes in Computer Science; Springer International Publishing: Cham, 2015; Vol. 9351; pp 234–241.

(8) Falk, T. et al. U-Net: deep learning for cell counting, detection, and morphometry. *Nature Methods* **2019**, *16*, 67–70.

(9) Abadi, M. et al. TensorFlow: Large-Scale Machine Learning on Heterogeneous Systems. 2015; `https://www.tensorflow.org/`.

(10) Abadi, M. et al. TensorFlow: A system for large-scale machine learning. Proceedings of the 12th USENIX Symposium on Operating Systems Design and Implementation (OSDI '16). Savannah, GA, USA, 2016.

(11) Pedregosa, F. et al. Scikit-learn: Machine Learning in Python. *Journal of Machine Learning Research* **2011**, *12*, 2825–2830.

(12) Zaharia, M.; Chen, A.; Davidson, A.; Ghodsi, A.; Hong, S. A.; Konwinski, A.; Murching, S.; Nykodym, T.; Ogilvie, P.; Parkhe, M.; Xie, F.; Zumar, C. Accelerating the Machine Learning Lifecycle with MLflow. *IEEE Data Engineering Bulletin* **2018**, *41*, 7.

(13) Chen, A. et al. Developments in MLflow: A System to Accelerate the Machine Learning Lifecycle. Proceedings of the Fourth International Workshop on Data Management for End-to-End Machine Learning. New York, NY, USA, 2020.

(14) Walsh, I. et al. DOME: recommendations for supervised machine learning validation in biology. *Nature Methods* **2021**, *18*, 1122–1127.

(15) Harris, C. R. et al. Array programming with NumPy. *Nature* **2020**, *585*, 357–362.

(16) McKinney, W. Data Structures for Statistical Computing in Python. Proceedings of the 9th Python in Science Conference. 2010; pp 56 – 61.

(17) The pandas development team, pandas-dev/pandas: Pandas. 2020; `https://doi.org/10.5281/zenodo.3509134`.

(18) Müller, P. Python multiple-tau algorithm. 2012; `https://pypi.python.org/pypi/multipletau/`.

(19) Wohland, T.; Rigler, R.; Vogel, H. The Standard Deviation in Fluorescence Correlation Spectroscopy. *Biophysical Journal* **2001**, *80*, 2987–2999.

(20) Schaetzel, K.; Peters, R. Noise on multiple-tau photon correlation data. Photon Correlation Spectroscopy: Multicomponent Systems. 1991; pp 109–115.

(21) Wahl, M.; Gregor, I.; Patting, M.; Enderlein, J. Fast calculation of fluorescence correlation data with asynchronous time-correlated single-photon counting. *Optics Express* **2003**, *11*, 3583.

(22) Waithe, D.; Clausen, M. P.; Sezgin, E.; Eggeling, C. FoCuS-point: software for STED fluorescence correlation and time-gated single photon counting. *Bioinformatics (Oxford, England)* **2016**, *32*, 958–960.

(23) Sezgin, E.; Schneider, F.; Galiani, S.; Urbančič, I.; Waithe, D.; Lagerholm, B. C.; Eggeling, C. Measuring nanoscale diffusion dynamics in cellular membranes with super-resolution STED–FCS. *Nature Protocols* **2019**,
